# Substantial differences in soil viral community composition within and among four Northern California habitats

**DOI:** 10.1101/2022.05.26.493654

**Authors:** Devyn M. Durham, Ella T. Sieradzki, Anneliek M. ter Horst, Christian Santos-Medellín, C. Winston A. Bess, Sara E. Geonczy, Joanne B. Emerson

**Affiliations:** Department of Plant Pathology, University of California Davis, CA, USA

## Abstract

Viruses contribute to food web dynamics and nutrient cycles in diverse ecosystems, yet the biogeographical patterns that underlie these viral dynamics are poorly understood, particularly in soil. Here, we identified trends in soil viral community composition in relation to habitat, moisture content, and physical distance. We generated 30 soil viromes from four distinct habitats (wetlands, grasslands, woodlands, and chaparral) by selectively capturing virus-sized particles prior to DNA extraction, and we recovered 3,432 unique viral ‘species’ (vOTUs). Viral communities differed significantly by soil moisture content, with viral richness generally higher in wet compared to dry soil habitats. However, vOTUs were rarely shared between samples, including replicates <10 m apart, suggesting that soil viruses may not disperse well and that future soil viral community sampling strategies may need to account for extreme community differences over small spatial scales. Of the 19% of vOTUs detected in more than one sample, 93% were from the same habitat and site, suggesting greater viral community similarity in closer proximity and under similar environmental conditions. Within-habitat differences indicate that extensive sampling would be required for rigorous cross-habitat comparisons, and results belie emerging paradigms of higher viral activity in wet soils and soil viral community spatial heterogeneity.

## Introduction

Viruses are abundant across Earth’s ecosystems, contributing to microbial dynamics and biogeochemical cycles, yet they remain understudied, particularly in terrestrial habitats [1, 2]. Soil viral abundance measurements vary substantially, ranging from nearly zero in dry deserts to over 10^9^ virus-like particles per gram in wetlands [3]. In the better studied oceans, viruses kill approximately 20% of microbial biomass daily, impacting nutrient and energy cycles [4], and recent work suggests that viruses may be similarly important in terrestrial ecosystems [3, 5–12]. For example, viruses have been suggested to affect carbon cycling in thawing permafrost peatlands by preying on methanogens and methanotrophs and by encoding glycoside hydrolases to break down complex carbon into simple sugars [2]. Soil viral activity has been demonstrated [2, 5, 13], and soil viral communities can be spatially structured [14–16]. Despite these emerging ecological patterns, comparisons of soil viral diversity within and across habitats are limited..

Here, we compared dsDNA viromes (viral-size fraction metagenomes) [14] from four distinct habitats (wetlands, grasslands, chaparral shrublands and oak woodlands) across five UC Davis Natural Reserves field sites within Northern California. We compared viral species (vOTU) richness, vOTU detection patterns, and viral community beta-diversity, according to habitat type, soil properties, and spatial distribution to better understand the fundamental relationships between soil viruses and the ecosystems that they inhabit.

## Results and Discussion

To compare soil viral community composition within and across terrestrial habitats on a regional scale, viromes were generated from 34 near-surface (top 15 cm) soil samples, with a total of 30 viromes included in downstream ecological analyses (see Supplementary Methods). The analyzed samples were collected from four distinct habitats (wetlands, grasslands, chaparral shrublands, and woodlands, each with 14, 7, 5, and 4 viromes, respectively) across five field sites (see Table S1 for sample locations and soil properties). Following quality filtering, the 30 viromes generated an average of 72,950,833 reads and 416 contigs ≥ 10 Kbp each (Table S2). Wetland viromes yielded more contigs ≥ 10 Kbp than viromes from other habitats (Table S2). We used VIBRANT to identify 3,490 viral contigs in our assemblies, which were clustered into 3,432 viral operational taxonomic units (vOTUs), defined as ≥ 10 Kbp viral contigs sharing ≥ 95% average nucleotide identity.

Multiple lines of evidence suggest that wetter soils harbored greater viral diversity than drier soils. We recovered the most vOTUs from wetlands, both in total (56% of all vOTUs were from wetlands) and per sample (on average, 307 vOTUs were recovered per wetland sample, compared to 116 from other habitats) (Fig. 1A). Unsurprisingly, wetlands had significantly greater moisture content than other habitats (Fig. 1B; ANOVA followed by Tukey multiple comparisons of means, p < 0.001), especially considering that samples were collected towards the end of the Mediterranean dry season. Although viral richness was qualitatively highest in wetlands, this was not statistically significant (ANOVA model richness ∼ habitat, p = 0.095). Nonparametric tests, which account for nonlinear correlations, revealed a significantly positive correlation between viral richness and soil moisture content (Spearman rho = 0.65, p < 0.001; Kendall tau = 0.43, p = 0.002). Viral community beta-diversity was also related to soil chemical properties overall (Mantel test, R^2^=0.43, p=0.009; Table S1), while distance between sites only accounted for 5% of the variation (Partial Mantel test, R^2^=0.38, p=0.009). Taken together, viral diversity was generally highest in wet soils.

**Figure 1.**
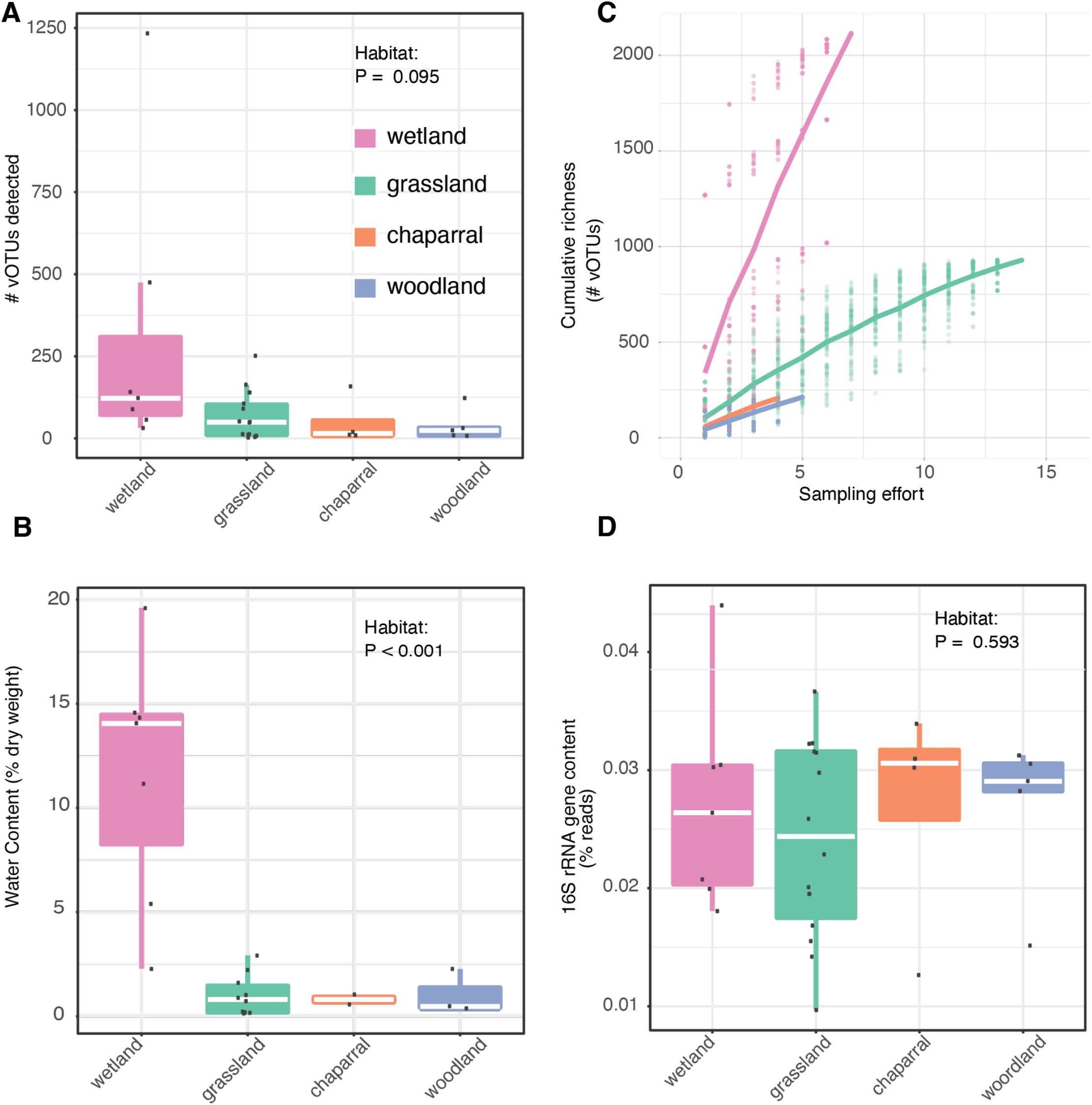
Evidence for higher viral richness in wetter soils. Comparisons between habitats of **(A)** Viral richness (number of identified vOTUs with coverage along at least 75% of the contig in a given virome), dots represent richness in individual samples, **(B)** Accumulation curves of cumulative vOTU richness as sampling effort increased, dots represent cumulative richness at each sampling effort across 100 permutations of sample order; the overlaid lines display the mean cumulative richness per habitat, **(C)** Water content, calculated as (wet weight – dry weight) divided by dry weight, and **(D)** Bacterial 16S rRNA gene content in the viromes, based on percent of viromic reads mapping to 16S rRNA reference genes. VIBRANT [19] was used to identify 3,490 viral contigs in our assemblies, and these viral contigs were clustered at 95% average nucleotide identity (ANI) into 3,432 viral operational taxonomic units (vOTUs). For A, B, D, raw data are plotted on top of the box plots, with white lines showing the median, boxes indicating 75% of the data, whiskers extending to 90%, and points beyond the whiskers indicating outliers.

We next wondered whether differences in sampling effort or bacterial content in the viromes could have produced the observed diversity patterns. For example, if viral richness had been captured more comprehensively in wetlands, and/or if bacterial content had been higher in viromes from other habitats, viral diversity could have appeared artificially higher in wetlands. A comparison of accumulation curves across habitats revealed the opposite: wetlands were likely to be the most under-sampled, in terms of true viral diversity (Fig. 1C). Given that relic DNA and small bacteria can pass through 0.22 μm filters, bacterial sequences are known to be present in viromes [14,17]. To compare bacterial content in viromes across habitats, 16S rRNA gene fragments were recovered from raw reads (Fig. 1D). The percentage of 16S rRNA gene sequences in each virome ranged from 0.01-0.044% (consistent with prior reports of 0.028% bacterial 16S rRNA gene content in similarly prepared viromes from agricultural soils [18]) and did not differ significantly by habitat (ANOVA, P = 0.595). Thus, viral diversity estimates did not seem to be disproportionately skewed by sampling effort or the presence of non-viral sequences in viromes.

Perhaps the most striking result from this study was the relative uniqueness of each soil viral community. The majority of vOTUs (81%) in this regional study were only detected in a single sample (Fig. 2A). Of the 666 vOTUs detected in multiple samples, 93% were detected in replicate samples from the same habitat and site (Fig. 2B). Samples the shortest distance apart (less than 1 Km) and from the same habitat and site (*i*.*e*., replicates) harbored the most similar viral communities (Fig. 2C). Within the same field site, viral communities were less similar between habitats, and paired samples from the same habitat at different, nearby sites (within 6 Km) did not share any vOTUs, suggesting substantial differences in viral communities over local distances both within and between habitats. At greater distances, community similarity generally decreased, even between samples from the same habitat (Fig. 2C). Still, 21 vOTUs were detected in multiple habitats across multiple sites (Fig. 2B), and some vOTUs were shared between the two farthest sites (109 Km apart, Fig. 2C), suggesting some degree of regional conservation of viral populations. Overall, results suggest substantial differences in soil viral community composition in the same habitat on the scale of meters, greater similarity of viral communities in close proximity and under similar environmental conditions, and a small number of vOTUs shared over regional distances.

**Figure 2.**
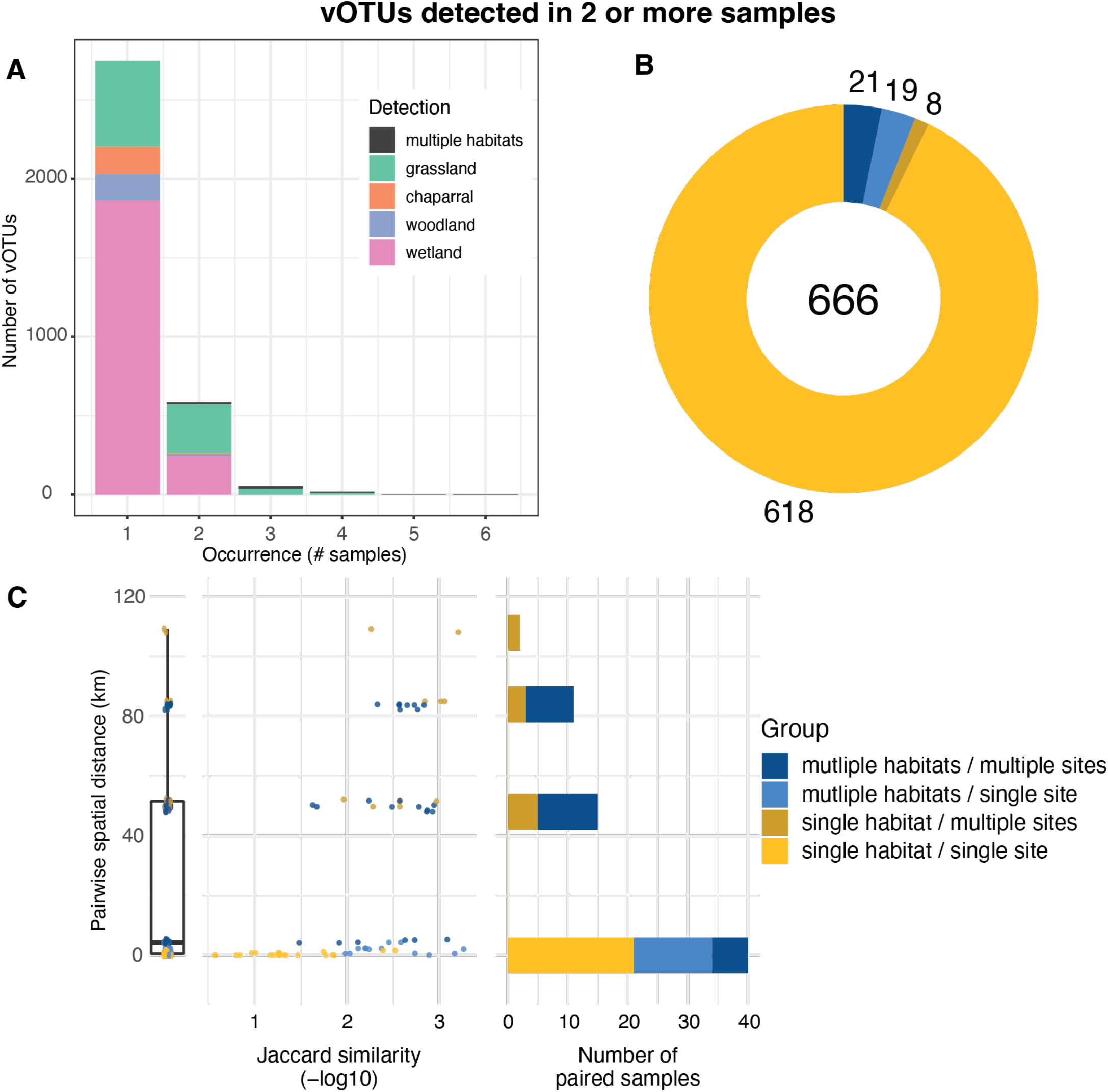
Substantial differences in vOTU detection and viral community composition between soil samples, both within and across habitats. **(A)** vOTU detection patterns, in terms of the number of samples in which each vOTU was detected (x-axis) and the number of vOTUs exhibiting a given detection pattern (y-axis), with stacked bar colors indicating the habitat(s) in which each vOTU was detected, **(B)** vOTUs detected in more than one sample, colored by their detection patterns across sites and habitats (legend to the right of panel C), numbers indicate total vOTUs (center) and vOTUs per detection pattern, and **(C)** Pairwise community compositional similarity (x-axis) by geographic distance (y-axis) between samples, colored according to detection patterns across habitats and sites (legend on the far right). Left graph: box plot of pairwise distances for all samples (condensed x-axis). Middle graph: Each point represents the - log10 Jaccard similarity between two samples along the x-axis, with lower values (left side of the x-axis) indicating greater viral community similarity. Right graph: Total number of sample pairs included at each geographic distance. Paired samples were only included if they shared at least one vOTU.

Although soil viral communities are woefully undersampled, making sweeping generalizations premature, results from this study and others converge on a picture of high local viral diversity, with communities differing substantially over space and by habitat, yet some regional and global conservation of viral ‘species’ (vOTUs). Viral community distance-decay relationships have been observed across an 18-m long agricultural field [14] and in a 200 m^2^ grassland [20], consistent with the meters-scale differences in viral community composition between replicates from the same habitat observed in this study. However, the proportion of vOTUs shared over meters varied across these studies, with many vOTUs shared across the agricultural field but most vOTUs not shared between samples ∼10 m apart in this study and in the grassland field, suggesting the potential for different spatial patterns in natural and managed soils, perhaps due to different amounts of mixing. Similarly substantial differences among viral communities on a regional scale were also identified in a study of grassland and peatland RNA viromes, which shared few viruses between samples [16]. However, ter Horst et al. showed that 4% of the vOTUs from a Minnesota, USA peatland were shared in other peatlands, often on different continents [8], consistent with the recovery of a small number of vOTUs shared over >100 Km distances here. Taken together, we propose that soil viral communities often display high heterogeneity within and among habitats, presumably due to a combination of host adaptations and microdiversity, dispersal limitation, and fluctuating environmental conditions over space and time.

## Acknowledgements

We thank the University of California, Davis Natural Reserves site directors and staff, particularly Shane Waddell, Catherine Koehler, Jeffrey Clary, Sarah Oktay, Suzanne Olyarnik, and Jacqueline Sones, for facilitating site access, showing us around the field sites, and providing logistical support for this work. Funding for this work was provided by new lab start-up to JBE from the University of California, Davis College of Agricultural and Environmental Sciences and Department of Plant Pathology, as well as by the U.S. Department of Energy (DOE), Office of Science, Office of Biological and Environmental Research (BER), Genomic Science Program, award number DE-SC0021198 (grant to JBE).

## Supplemental Methods

### Experimental design adjustments on account of COVID-19 and wildfire

The sample set analyzed in this study, collected in November 2019, was meant to be the first of several time points, with plans for periodic sampling of the same sites in 2020-2021.

However, COVID-19 prevented sample collection throughout much of 2020, and then the LNU Complex Fires completely burned 2 and partially burned 1 of our 5 field sites in August 2020. Thus, the decision was made to analyze this November 2019 dataset on its own.

### Site description and field sampling

Soil samples were collected from woodland, chaparral, grassland, coastal bluff, and wetland habits at five UC Davis Natural Reserves sites in Northern California, USA: Bodega Bay (BB), Jepson Prairie (JP), McLaughlin (ML), Quail Ridge (QR), and Stebbins Cold Canyon (SCC). From November 14 through November 16, 2019, 34 samples were collected from 17 plots within the five sites. We sampled at least two distinct habitats from each site, and two samples (replicates) were retrieved from each plot, with each plot consisting of one contiguous habitat. Inter-sample distance between replicates in each plot ranged from 6.5-74.2 m and was meant to capture a large area within one instance of the habitat (*i*.*e*., no distinct habitat types were permitted between replicate samples), considering available habitat space and accessibility. The number of samples per site and habitat, along with GPS coordinates (Table S1) and distance between sites, appear in Tables S3 and S4, and soil chemistry appears in Table S1.

For each sample, three 7.6 cm diameter soil cores were collected from the 0-15 cm depth range (excluding aboveground biomass) with a slide-hammer corer, approximately 15 cm apart from each other. The three fresh cores for each sample were homogenized and transferred into sterile bags on site, kept on ice for 1-2 days in the field, and then passed through an 8 mm mesh sieve within 3 days of sample collection and then stored at -80 °C prior to processing for DNA extraction and soil properties.

### Soil viral purification and DNA extraction

Viral particle purification and DNA extraction proceeded similarly to the Santos-Medellin et al. [1] protocol (which was largely derived from Goller et al. [2]), with slight modifications. Briefly, for each sample, a suspended soil solution was prepared by adding 9 mL of PPBS Buffer (2% bovine serum albumin, 10% phosphate-buffered saline, 1% potassium citrate, and 150 mM MgSO4) to 10 g of soil [2]. To elute virions, soil suspensions were shaken by inversion and placed in an orbital shaker for 20 min at 300 RPM and 4 °C, then centrifuged (Sorvall RC5C, Thermo-Fisher Scientific, Waltham, MA, USA) for 10 min at 10,000 x g and 4 °C. Supernatant was briefly stored at 4 °C while the soil pellet was resuspended with 9 mL of PPBS Buffer, and the entire process of mixing, shaking, and centrifugation was repeated two more times, resulting in a total of ∼27 mL of supernatant per soil sample. To pellet and remove remaining soil, supernatant was centrifuged for 8 min at 5,000 x g and 4 °C, with supernatant transferred, excluding the pellet. This step was repeated twice. The pooled supernatants were filtered through a 0.22 μm polyether sulfone filter and centrifuged (Optima LE-80K ultracentrifuge, Beckman Coulter, Brea, CA, USA; 50.2 Ti rotor) for 2 hours 25 minutes at 32,000 g and 4 °C. The supernatant was removed and the pellet containing viral particles was resuspended in 100 μL of ultrapure water. DNase was not used here, as samples were frozen, which may compromise the integrity of viral capsids [3, 4]. However, ecological trends from viromes can still be detected without DNase treatment [3, 4].

For DNA extraction, 250 μL of phenol:chloroform:isoamyl alcohol (pH 8) was added to each 100 μL sample. The mixture was vortexed and then incubated on ice for 3 cycles of 1 min on and 5 min off intervals. Samples were then centrifuged for 5 min at 14 000 x g and the supernatant was transferred to a Phase Lock Gel™ tube, and then centrifuged for 5 min at 14,000 x g. The viral DNA supernatant (100 μL) was mixed with 25 μL of 3 M sodium acetate, 1.5 μL of Glycoblue™ Coprecipitant, and 250 μL isopropanol. The mixture was vortexed and then incubated at -80 °C for 20 min. After incubation, samples were centrifuged for 20 min at 14,000 x g. Supernatant was removed and the viral pellet was washed with 500 μL of 70 % ethanol and centrifuged for 5 min at 14,000 x g. The ethanol was removed and the samples were centrifuged for another minute, then left to dry for 5 min. The viral DNA pellet was resuspended with 100 μL of non-DEPC-treated ultrapure water. DNA yields were quantified using an Invitrogen Qubit 4 Fluorometer with 1X High Sensitivity DNA assay (Thermo Fisher Scientific, Inc., Waltham, MA, USA). Of the 34 samples from which viral DNA was extracted, one replicate from a BB wetland site (sample ID: BB_3_2) and one replicate from a QR woodland site (sample ID: QR_2_1) yielded insufficient DNA and were thus excluded, resulting in 32 samples remaining for library construction and sequencing.

### Library construction and sequencing

Libraries from the 32 viromes were constructed with the KAPA Roche kit, as described in Santos-Medellin et al. [1], pooled in equimolar concentrations, and sequenced to a target depth of 10 Gbp per virome on the Illumina NovaSeq platform at the UC Davis DNA Technologies Core.

### Bioinformatics

Raw sequencing reads were quality-filtered and trimmed using Trimmomatic-0.39, as in Roux et al. [5, 6]. Removal of PhiX147 sequences was performed with BBDUK from the BBMap package (version: BBMap-38.87) [7], followed by assembly with MEGAHIT-1.2.9 in meta-large mode [8, 9]. Predicted viral contigs were identified using VIBRANT-1.2.1 with default settings [10]. dRep version 2.0.0 [11] was used for clustering sequences at 95% average nucleotide identity across 85% of the length of the shorter contig, resulting in a reference set of 4,712 viral population sequences (vOTUs). Reads from the 32 viromes were mapped to these reference vOTU sequences with Bowtie 2-2.4.2 [12], using sensitive mode, and the vOTU read-mapped coverage table was created with BamM-1.7.3 [13]. Where vOTU breadth (contig length fraction with at least 1x coverage depth) was below 75%, coverage was set to zero [8]. Two viromes (sample IDs: BB_6_1 and BB_6_2) were excluded from downstream comparative analyses at this point because they were the only samples from a coastal bluff habitat across the dataset, thus this habitat was deemed insufficiently sampled for cross-habitat comparisons. The final dataset for comparative analyses consisted of 30 viromes.

16S rRNA gene abundance recovery from viromes was performed by identifying reads that map to this gene with SortMeRNA v4.2.0 [15] against the SILVA database [16], as in [4,5].

### Soil properties

Soil was weighed upon arrival to the lab (Wet Weight, WW), allowed to dry completely in a biological hood, and weighed again (Dry Weight, DW). Soil water content was calculated as (WW-DW)/DW.

All other soil properties appearing in Table S1 were analyzed by Ward Laboratories (Kearney, NE, USA). Phosphorus and sulfate were extracted with Mehlich-3 buffer. Iron, manganese, copper, and zinc were extracted with DTPA buffer. Sodium, potassium, calcium, and magnesium were extracted with ammonium-acetate. Soil organic matter was calculated by percentage loss on ignition (LOI). Soil pH and soluble salts were measured using a 1:1 soil:water suspension. Nitrate was extracted with KCl.

### Data analysis

Data analysis was performed in R on the vOTU read-mapped coverage table generated previously (see *Bioinformatics*). When calculating community similarity (beta-diversity), we used Jaccard similarity, as opposed to Bray-Curtis similarity, as there were so few vOTUs shared between samples. Similarity matrices were calculated using vegan [17]. Richness of vOTUs detected per site and per habitat was calculated using tidyverse [18] and plotted with cowplot [19] and ggplot2 [20]. Comparisons of vOTU richness between sites were performed with rstatix [21], using the Games-Howell test to account for uneven sample size between habitats. Correlations between water content and viral richness, ANOVA comparison of water content between environments, and Tukey post hoc tests were calculated using the R stats package. The correlation analysis was performed under the hypothesis that higher moisture leads to higher richness and was therefore one-sided (alternative=greater). A Mantel test comparing viral community composition to soil chemistry was performed by creating Jaccard distance matrices (vegdist) and comparing them (mantel, 100 permutations; mantel.partial, 100 permutations) with functions from the Vegan R package [18].

### Data availability

The datasets generated and/or analyzed in this study have been submitted to the NCBI sequence read archive (SRA) under BioProject number PRJNA831438 and will become available upon publication. The vOTU fasta sequences are available on https://github.com/ellasiera/Nat_res_vOTUs.

## Supplementary Tables

Table S1. Soil chemistry characteristics of all samples.

**Table S2.**
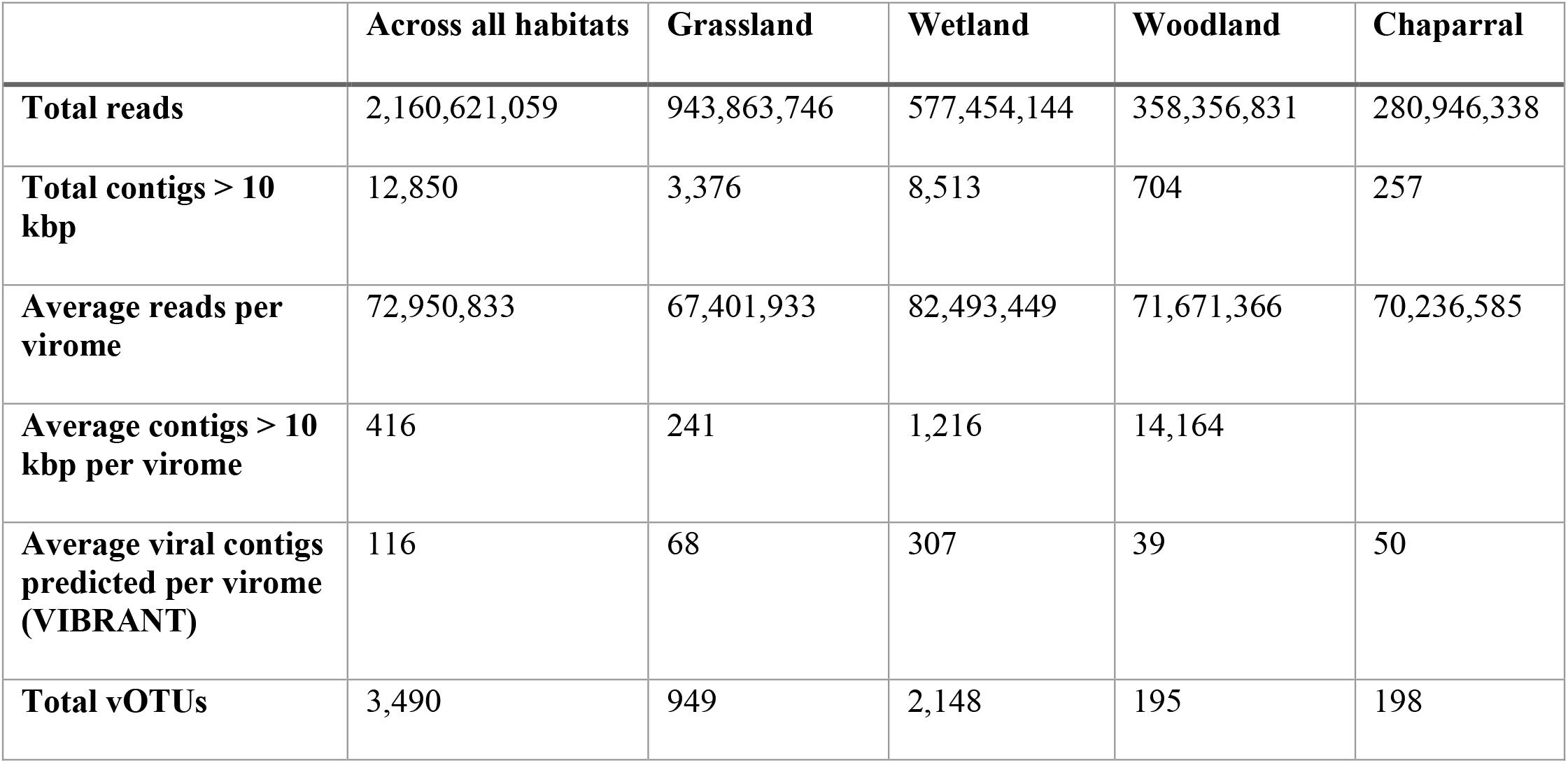
Summary of assembly statistics including detection of viral genomes per habitat

**Table S3.**
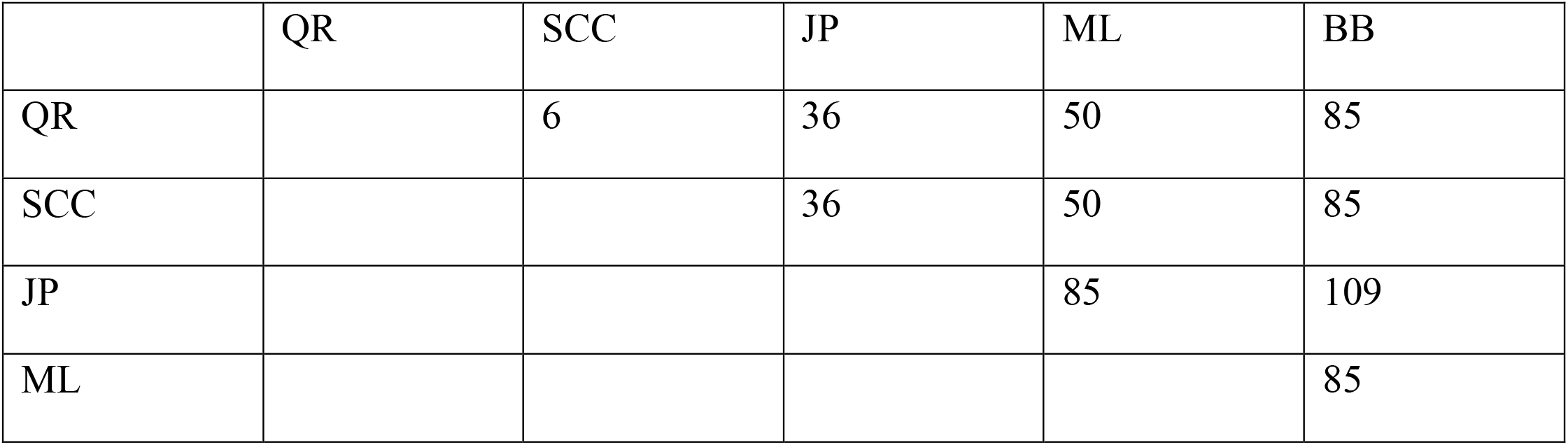
Distance (km) between UC Davis Natural Reserves sites sampled

**Table S4.**
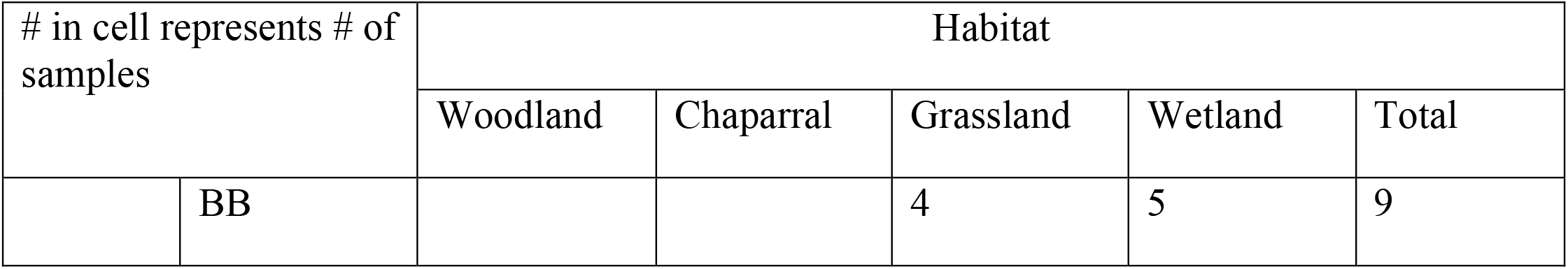
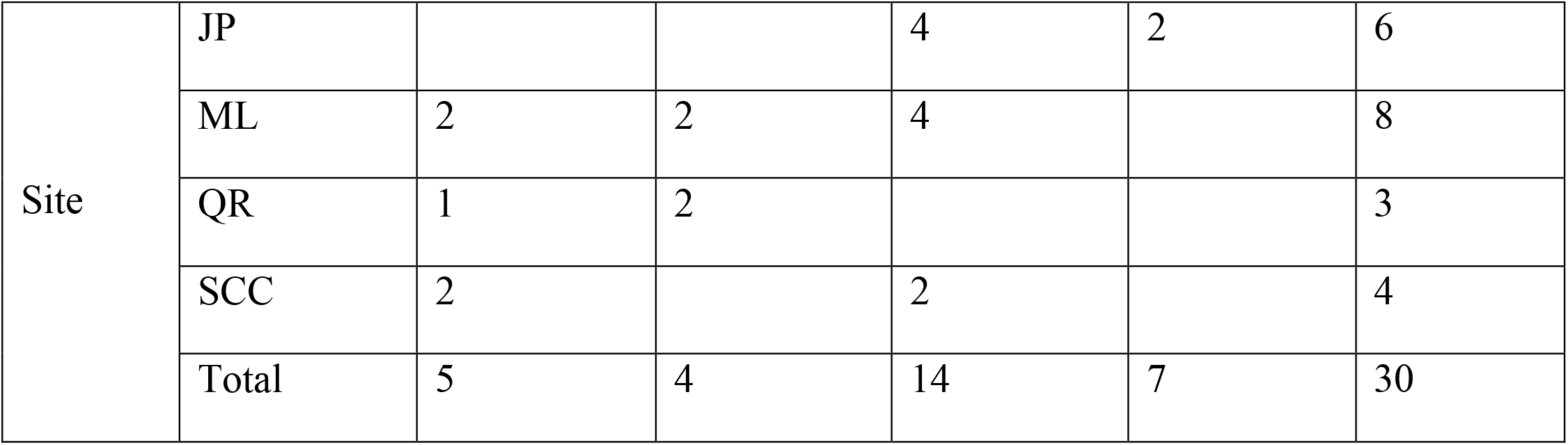
Number of samples analyzed by habitat and site

| # in cell represents # of samples |  | Habitat |  |  |  |  |
| --- | --- | --- | --- | --- | --- | --- |
|  |  | Woodland | Chaparral | Grassland | Wetland | Total |
|  | BB |  |  | 4 | 5 | 9 |
| Site | JP |  |  | 4 | 2 | 6 |
|  | ML | 2 | 2 | 4 |  | 8 |
|  | QR | 1 | 2 |  |  | 3 |
|  | SCC | 2 |  | 2 |  | 4 |
|  | Total | 5 | 4 | 14 | 7 | 30 |

