## Supplemental methods for "Substantial differences in soil viral community composition within and among four Northern California habitats"

*Bioinformatics*

Raw sequencing reads were quality-filtered and trimmed using Trimmomatic-0.39, as in Roux et al. [[5, 6]](https://paperpile.com/c/hsgGpF/gkZV+PMYh). Removal of PhiX147 sequences was performed with BBDUK from the BBMap package (version: BBMap-38.87) [[7]](https://paperpile.com/c/hsgGpF/OtJt), followed by assembly with MEGAHIT-1.2.9 in meta-large mode [[8, 9]](https://paperpile.com/c/hsgGpF/a23t+0hJh). Predicted viral contigs were identified using VIBRANT-1.2.1 with default settings [[10]](https://paperpile.com/c/hsgGpF/15oU). dRep version 2.0.0 [[11]](https://paperpile.com/c/hsgGpF/etkR) was used for clustering sequences at 95% average nucleotide identity across 85% of the length of the shorter contig, resulting in a reference set of 4,712 viral population sequences (vOTUs). Reads from the 32 viromes were mapped to these reference vOTU sequences with Bowtie 2-2.4.2 [[12]](https://paperpile.com/c/hsgGpF/r3L4), using sensitive mode, and the vOTU read-mapped coverage table was created with BamM-1.7.3 [[13]](https://paperpile.com/c/hsgGpF/Xi2g). Where vOTU breadth (contig length fraction with at least 1x coverage depth) was below 75%, coverage was set to zero [[8]](https://paperpile.com/c/hsgGpF/a23t). Two viromes (sample IDs: BB_6_1 and BB_6_2) were excluded from downstream comparative analyses at this point because they were the only samples from a coastal bluff habitat across the dataset, thus this habitat was deemed insufficiently sampled for cross-habitat comparisons. The final dataset for comparative analyses consisted of 30 viromes.

16S rRNA gene abundance recovery from viromes was performed by identifying reads that map to this gene with SortMeRNA v4.2.0 [[15]](https://paperpile.com/c/MnOQcM/s2Ub) against the SILVA database [[16]](https://paperpile.com/c/MnOQcM/uJyr), as in [4,5].

*Soil properties*

Soil was weighed upon arrival to the lab (Wet Weight, WW), allowed to dry completely in a biological hood, and weighed again (Dry Weight, DW). Soil water content was calculated as (WW-DW)/DW.

All other soil properties appearing in Table S1 were analyzed by Ward Laboratories (Kearney, NE, USA). Phosphorus and sulfate were extracted with Mehlich-3 buffer. Iron, manganese, copper, and zinc were extracted with DTPA buffer. Sodium, potassium, calcium, and magnesium were extracted with ammonium-acetate. Soil organic matter was calculated by percentage loss on ignition (LOI). Soil pH and soluble salts were measured using a 1:1 soil:water suspension. Nitrate was extracted with KCl.

*Data analysis*

Data analysis was performed in R on the vOTU read-mapped coverage table generated previously (see *Bioinformatics*). When calculating community similarity (beta-diversity), we used Jaccard similarity, as opposed to Bray-Curtis similarity, as there were so few vOTUs shared between samples. Similarity matrices were calculated using vegan [[17]](https://paperpile.com/c/MnOQcM/2O0S). Richness of vOTUs detected per site and per habitat was calculated using tidyverse [[18]](https://paperpile.com/c/MnOQcM/frO3) and plotted with cowplot [[19]](https://paperpile.com/c/MnOQcM/MpGq) and ggplot2 [[20]](https://paperpile.com/c/MnOQcM/Ud8O). Comparisons of vOTU richness between sites were performed with rstatix [[21]](https://paperpile.com/c/MnOQcM/uAaq), using the Games-Howell test to account for uneven sample size between habitats. Correlations between water content and viral richness, ANOVA comparison of water content between environments, and Tukey post hoc tests were calculated using the R stats package. The correlation analysis was performed under the hypothesis that higher moisture leads to higher richness and was therefore one-sided (alternative=greater). A Mantel test comparing viral community composition to soil chemistry was performed by creating Jaccard distance matrices (vegdist) and comparing them (mantel, 100 permutations; mantel.partial, 100 permutations) with functions from the Vegan R package [[18]](https://paperpile.com/c/MnOQcM/2O0S).

*Data availability*

The datasets generated and/or analyzed in this study have been submitted to the NCBI sequence read archive (SRA) under BioProject number PRJNA831438 and will become available upon publication. The vOTU fasta sequences are available on https://github.com/ellasiera/Nat_res_vOTUs.
