## Supplemental table S2 for "Substantial differences in soil viral community composition within and among four Northern California habitats"

Table S2. Summary of assembly statistics including detection of viral genomes per habitat

|  | Across all habitats | Grassland | Wetland | Woodland | Chaparral |
| --- | --- | --- | --- | --- | --- |
| Total reads | 2,160,621,059 | 943,863,746 | 577,454,144 | 358,356,831 | 280,946,338 |
| Total contigs > 10 kbp | 12,850 | 3,376 | 8,513 | 704 | 257 |
| Average reads per virome | 72,950,833 | 67,401,933 | 82,493,449 | 71,671,366 | 70,236,585 |
| Average contigs > 10 kbp per virome | 416 | 241 | 1,216 | 14,164 |  |
| Average viral contigs predicted per virome (VIBRANT) | 116 | 68 | 307 | 39 | 50 |
| Total vOTUs | 3,490 | 949 | 2,148 | 195 | 198 |
