## Supplemental table S3 for "Substantial differences in soil viral community composition within and among four Northern California habitats"

Table S3. Distance (km) between UC Davis Natural Reserves sites sampled

|  | QR | SCC | JP | ML | BB |
| --- | --- | --- | --- | --- | --- |
| QR |  | 6 | 36 | 50 | 85 |
| SCC |  |  | 36 | 50 | 85 |
| JP |  |  |  | 85 | 109 |
| ML |  |  |  |  | 85 |
