## Supplemental table S4 for "Substantial differences in soil viral community composition within and among four Northern California habitats"

Table S4. Number of samples analyzed by habitat and site

| # in cell represents # of samples | | Habitat | | | | |
| --- | --- | --- | --- | --- | --- | --- |
|  |  | Woodland | Chaparral | Grassland | Wetland | Total |
| Site | BB |  |  | 4 | 5 | 9 |
|  | JP |  |  | 4 | 2 | 6 |
|  | ML | 2 | 2 | 4 |  | 8 |
|  | QR | 1 | 2 |  |  | 3 |
|  | SCC | 2 |  | 2 |  | 4 |
|  | Total | 5 | 4 | 14 | 7 | 30 |
